# Chronic Social Defeat Stress Gives Rise to Social Avoidance Through Fear Learning

**DOI:** 10.1101/2024.06.18.599597

**Authors:** Jinah Lee, Antonio Aubry, Sadiyah Hanif, Itamar Grunfeld, Ekaterina Likhtik, Nesha S. Burghardt

## Abstract

Chronic social defeat stress (CSDS), a widely used rodent model of stress, reliably leads to decreased social interaction in stress susceptible animals. Here, we investigate a role for fear learning in this response using 129Sv/Ev mice, a strain that is more vulnerable to CSDS than the commonly used C57BL/6 strain. We first demonstrate that defeated 129Sv/Ev mice avoid a CD-1 mouse, but not a conspecific, indicating that motivation to socialize is intact in this strain. CD-1 avoidance is characterized by approach behavior that results in running in the opposite direction, activity that is consistent with a threat response. We next test whether CD-1 avoidance is subject to the same behavioral changes found in traditional models of Pavlovian fear conditioning. We find that associative learning occurs across 10 days CSDS, with defeated mice learning to associate the color of the CD-1 coat with threat. This leads to the gradual acquisition of avoidance behavior, a conditioned response that can be extinguished with 7 days of repeated social interaction testing (5 tests/day). Pairing a CD-1 with a tone leads to second-order conditioning, resulting in avoidance of an enclosure without a social target. Finally, we show that social interaction with a conspecific is a highly variable response in defeated mice that may reflect individual differences in generalization of fear to other social targets. Our data indicate that fear conditioning to a social target is a key component of CSDS, implicating the involvement of fear circuits in social avoidance.

## 1. Introduction

Chronic stress is a major risk factor for developing mood and anxiety disorders [1–3]. As such, preclinical animal models of psychiatric disorders commonly involve exposing rodents to chronic stressors, with chronic social defeat stress (CSDS) being among the most popular [4, 5]. In CSDS, repeated exposure to an aggressive CD-1 mouse reliably leads to enduring deficits in social interaction with a novel CD-1 in stress susceptible mice. In addition, CSDS perturbs social conditioned place preference [6] and reduces preference for other natural rewards, such as sucrose [7]. The finding that defeated mice also avoid juvenile [6] and adult mice of the same strain (i.e., conspecifics) [8–10] further suggests that CSDS decreases motivation to socialize. Given that social avoidance is often found in depressed humans, this is often interpreted as depressive-like behavior.

Notably, social avoidance is also a maladaptive coping strategy commonly observed in individuals with social anxiety disorder and post-traumatic stress disorder. Social avoidance reduces the risk of confronting a known stressor and can impair daily functioning and prolong anxiety by preventing exposure to experiences that promote extinction [11]. CSDS-induced social avoidance may therefore also reflect clinically relevant fear-based behavior that prevents the benefits of exposure therapy. However, few studies have investigated the role of fear learning in CSDS-induced social avoidance.

We hypothesize that fear conditioning occurs during each social defeat session, such that the CD-1 aggressor serves as a conditioned stimulus (CS) and the CD-1 attack serves as an unconditioned stimulus (US), leading to the formation of a fear memory that is retrieved during the social interaction test. Here, we evaluate this hypothesis by testing whether social avoidance is subject to the same behavioral changes found in traditional models of Pavlovian fear conditioning. We conducted all experiments in 129Sv/Ev mice, which we have previously shown to be more susceptible to CSDS than the more commonly used C57Bl/6 mouse strain [12]. We find that unlike C57Bl/6 mice, 129Sv/Ev mice only avoid CD-1 mice and not a conspecific, indicating that motivation to socialize is intact in this strain. We demonstrate that avoidance of the CD-1 gradually emerges across days of defeat as fear learning is acquired, during which defeated mice learn to associate the color of the CD-1 coat with threat. Like other forms of fear conditioning, we also show that avoidance of the CD-1 is subject to extinction and second-order fear conditioning. Finally, our results demonstrate that following CSDS, social interaction with a conspecific is a highly variable response that may be associated with individual differences in generalization of fear. Collectively, our results indicate that CSDS in 129Sv/Ev mice may be useful for studying the acquisition and retrieval of fear memories involving a social cue, a topic that is particularly relevant given the high rate of in-person and cyberbullying experienced by children and adults [13].

## 2. Materials and Methods

### 2.1 Animals

Eight-week-old male 129Sv/Ev mice and 12-week-old male Swiss webster mice (Taconic Biosciences, Germantown, NY) were group housed (2-4 mice/cage) upon arrival with mice of the same strain. Retired male CD-1 breeders (Charles River Laboratories, Wilmington, MA) were individually housed upon arrival. Mice were maintained on a 12-hour light-dark cycle (08:00-20:00) with free access to food and water and were given a minimum of one week to acclimate to the colony room before experiments began. Experiments were conducted in accordance with NIH guidelines and were approved by the Institutional Animal Care and Use Committee of Hunter College.

### 2.2 Chronic social defeat stress

#### Screening

CD-1 mice were individually housed in large plastic cages (30.8cm x 30.8cm x 14.29cm) (Thoren Caging Systems, Hazleton, PA) modified to accommodate a plexiglass divider at least 3 days before they were screened for aggressive behavior. Screening lasted 2-4 days, during which a 129Sv/Ev screener mouse was placed inside the home cage of the CD-1 and latency to attack was recorded. CD-1 mice were included in the experiment if they attacked the screener within 30 seconds for 2 consecutive days. CD-1 mice remained in these large cages until the experiment ended.

#### Chronic social defeat stress (CSDS)

CSDS was performed as previously described [4, 12], unless otherwise specified. Experimental mice were placed in the home cage of a CD-1 aggressor where they were allowed to interact for 5 minutes. Each session was monitored closely by the experimenter and terminated early if bite wounds resulted in bleeding. Mice were then separated overnight by a clear perforated plexiglass divider placed inside the aggressors’ home cage. This prevented further physical contact but allowed for continuous exposure to stressful sensory cues. This procedure was repeated for 10 consecutive days, with a novel CD-1 aggressor each day. Non-defeated control mice were pair housed in the same large cages, but were never exposed to a CD-1 aggressor. Instead, they interacted with a novel mouse of same strain each day for 5 minutes before being separated overnight by a perforated divider. After the final defeat session, all mice were individually housed in standard mouse cages.

### 2.3 Chronic restraint stress

The 12-hour light-dark cycle was adjusted for mice in the restraint stress and non-stress control groups, with lights on at 12:00. Mice were pair housed with mice of the same group in the same large cages used for CSDS, with one mouse on either side of the plexiglass divider. Each day, mice in the stress group were removed from the colony room at variable times of the dark cycle, weighed, and then individually restrained for 1 hour in plastic DecapiCones (Braintree Scientific, Braintree, MA) that were secured with a binder clip. The lights were turned on throughout the duration of restraint stress. At the same time, mice in the control group remained in the colony room where they were weighed and handled for 5 minutes in the dark by an experimenter wearing a red headlamp. Like CSDS, mice experienced restraint stress for 10 consecutive days.

### 2.4 Social interaction test

The social interaction test was conducted as previously described [12, 14]. Mice were placed in the center of an open field arena (25cm x 48cm) containing two identical wire-mesh enclosures located in diagonal corners of the box. One enclosure contained a novel social target and the other was empty. For experiments in which social interaction was tested repeatedly in the same mice, a novel social target was used in each test, unless otherwise specified. When mice were tested with more than one type of social target (e.g., CD-1, 129Sv/Ev, and Swiss Webster), the order of testing was counterbalanced. Test sessions lasted 5 minutes and were videotaped for later scoring. An observer blind to group quantified the number of seconds mice spent within each interaction zone, defined as the square region (12.5 cm× 12.5 cm) surrounding each enclosure. A defeat index (DI) was calculated by dividing the difference in time spent with each enclosure (social target enclosure – empty enclosure) by total time with both enclosures.

#### Acquisition of Social Avoidance

In a subset of mice, interaction with a novel CD-1 was tested the day before exposure to chronic stress and again after 1, 4, 7, and 10 days of CSDS or restraint stress. When social interaction testing and stress exposure occurred on the same day, social interaction was tested immediately *before* CSDS and an average of 4-hours *before* restraint stress. On each day of CSDS, mice were weighed immediately before defeat.

#### Interaction with CD-1 Scent

To evaluate whether exposure to the scent of a CD-1 elicits avoidance behavior, mice were placed in the same arena used for social interaction testing, but one enclosure was filled with bedding from the cage of a novel CD-1 mouse, including feces. The other enclosure remained empty. Time spent with each enclosure was quantified, as described above.

### 2.5 Extinction of Social Avoidance

One day after the last day of CSDS, social interaction was tested 5 times per day (inter-trial interval 1-hr) for 7 consecutive days. Half of the mice were only tested with a CD-1 mouse and half were only tested with a 129Sv/Ev mouse as the social target. Each social interaction test lasted 5 minutes. The CD-1 social target was novel on trials 1-25 and the 129Sv/Ev social target was novel on trials 1-22. After that, mice were exposed to a social target they had seen before, but they were only exposed to the same mouse a maximum of 2 times.

### 2.6 Second-Order Fear Conditioning

#### Conditioning

During each day of CSDS, the CD-1 aggressor was removed from its home cage and placed in a holding cage. A mouse in the defeated group was then placed in the empty home cage of the CD-1 for one minute before a tone (4kHz, 70dB, 30 seconds) played from an external speaker placed ∼3-inches away from the cage. During the last 10 seconds of the tone, the CD-1 was returned to its home cage (tone-CD1 pairing), where it was allowed to interact with the experimental mouse for 5 minutes. The mice were then separated overnight by a clear perforated plexiglass divider within the aggressors’ home cage. This procedure was repeated for 10 consecutive days with a novel CD-1 each day. Mice in the control group were housed in the same CSDS control conditions described above, but heard the tone (4kHz, 70dB, 30 seconds) once/day in their home cage while separated by the plexiglass divider. Therefore, controls were exposed to the tone as often as defeated mice, but never associated the tone with social interaction. An observer blind to group scored videos of each conditioning day and quantified seconds freezing to the context during the 60-second period before tone onset (contextual freezing) and freezing during the first 20-seconds of the tone presentation, prior to the arrival of the CD-1 (tone freezing). Freezing was defined as the cessation of all movement unrelated to respiration, for at least 1 second.

#### Tone Testing

Defeated and non-defeated controls were placed in the same arena used for the social interaction test, but both enclosures were empty. When a mouse entered the interaction zone of one enclosure (tone-paired enclosure), the same tone used during conditioning (4kHz, 70dB) played continuously until the mouse left the area. Time spent within the interaction zone of each enclosure was scored by an experimenter blind to group. The interaction zone was defined as the square region (12.5 cm × 12.5 cm) surrounding each enclosure, as described above.

### 2.7 Differential fear conditioning

One day after the last social interaction test, mice underwent a 5-day Pavlovian differential fear conditioning paradigm, as previously described [15, 16]. This included 1 day of habituation, 3 days of discrimination training and 1 day of retrieval testing. Mice were given 1-2 hours to acclimate to a room adjacent to the fear conditioning room before each session began. All behavior was recorded with an overhead camera and scored offline by an observer blind to the group.

#### Habituation

Mice were habituated to handling (5 minutes), the training context and then the testing context, with at least 1 hour between context exposures. The training context contained a metal grid floor (Med Associates, Inc., St. Albans, VT), 40 lux overhead lighting, and was cleaned between animals with ethanol. Mice were exposed to two separate pure tones (2kHz and 8kHz) in the training context. Tones were presented in pseudorandom order as 30 pips (50ms) amplitude modulated with a linear 25ms increase and 25ms decrease, once/second for 30 total seconds. Tones were presented with a variable inter-trial interval (ITI) of 60, 90, or 120 seconds. Then mice were habituated for 8 minutes to the testing context, which consisted of a wooden enclosure (45 cm length x 13 cm width x 20 cm height) containing a smooth paper floor that was changed between animals and ∼70 lux lighting. Tones were not played during habituation to the testing context.

#### Discrimination Training and Testing

During training, mice were placed in the training context where they were exposed to 6 tones (2kHz, CS+) that co-terminated with a foot shock (0.5mA, 1 second) and 6 tones that were explicitly unpaired with shock (8kHz, CS-; no shock). During retrieval testing, mice were placed in the testing context where they were exposed to 6 CS+ tones and 6 CS-tones and were never shocked. Tones were always presented in pseudorandom order with the same variable ITI used during habituation. An observer blind to group quantified time spent freezing during each CS tone presented on the day of test, with freezing defined as cessation of movement, as described above.

### 2.8 Statistical Analyses

Data were analyzed with a Student’s t-test for independent samples, Welch’s t-test, Pearson correlation, and two-way ANOVA using GraphPad Prism software (GraphPad, San Diego, CA). Significance levels were set at p<0.05 and interactions were followed by Tukey’s HSD or Uncorrected Fisher’s LSD post hoc tests.

## 3. Results

### 3.1 Defeated 129Sv/Ev mice avoid an aggressive CD-1, but not a mouse of the same strain

Previous studies conducted in C57BL/6 mice indicate that CSDS induces social avoidance of both a CD-1 mouse and a conspecific[6, 8–10]. We tested whether this is also true in 129Sv/Ev mice by exposing them to 10 days of CSDS and quantifying time spent interacting with either a novel CD-1 or a novel 129Sv/Ev mouse one day later (Fig. 1A). A two-way ANOVA on interaction time revealed a significant stress × social target interaction (F_(1,55) =_ 12.68, p < 0.001). Defeated mice spent less time with the CD-1 than control mice (p < 0.001), but approximately the same amount of time with the 129Sv/Ev mouse as controls (Fig. 1B, 1C). Within group comparisons revealed that defeated mice spent more time with the social target when it was a novel conspecific than a CD-1 (p < 0.0001), indicating that motivation to socialize was intact. This was confirmed when defeat-index (DI) scores were compared across groups (stress x social target interaction, F_(1,55)_ = 16.49, p < 0.001), which takes into account time with the empty enclosure during the social interaction test. We found that DI scores were lower in defeated mice tested with a CD-1 than controls tested with a CD-1 (p<0.0001) and similarly high in both groups when the 129Sv/Ev mouse was the social target. DI scores in defeated mice were also higher when the social target was the 129Sv/Ev than the CD-1 (p < 0.0001) (Fig. 1D). To evaluate the possibility that avoidance of the CD-1 reflects a threat response, we quantified the number of times mice entered the center of the arena to approach the CD-1 and the number of times this was immediately followed by running in the opposite direction (i.e., avoidance). We found that when defeated mice approached the CD-1, they ran away ∼97% of the time, which is significantly higher than control mice, which ran away from the CD-1 ∼42% of the time (p<0.001). A two-way ANOVA on percentage of approaches resulting in avoidance revealed a significant stress x social target interaction (F_(1,35)_ = 5.42, p<0.05), with defeated mice running away more often when they approached the CD-1 than the 129Sv/Ev (p<0.01) (Fig. 1E). Such active avoidance specifically of the CD-1 suggests that defeated mice may have learned to fear this social target.

**Figure 1.**
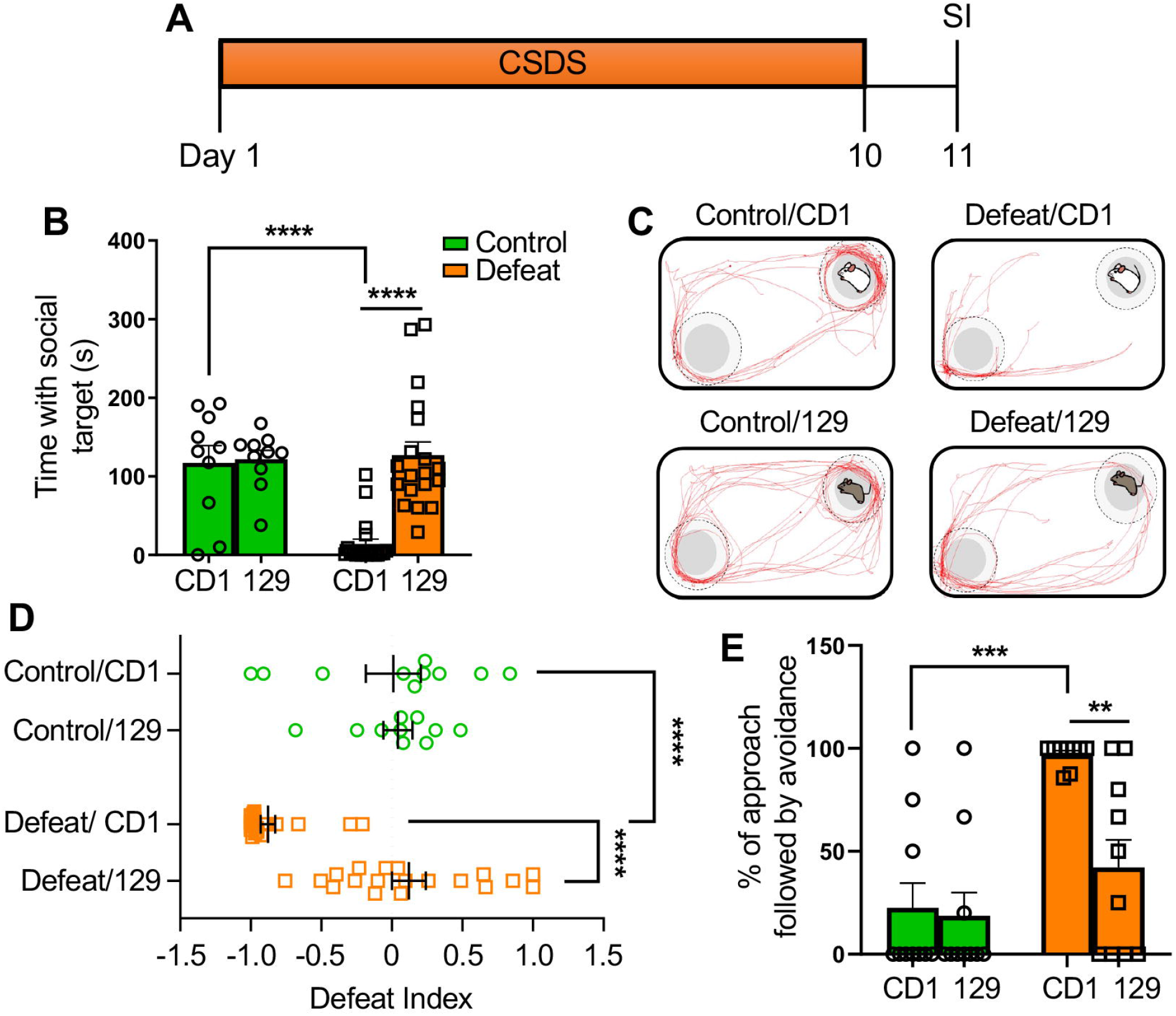
Defeated 129Sv/Ev mice do not avoid mice of the same strain. **(A)** Schematic of behavioral procedures. Social interaction was tested 1 day after the last defeat session with either a novel CD-1 or a novel conspecific as the social target. **(B)** Time spent with the enclosure containing each social target. Defeated mice avoided the CD-1, but spent as much time with the 129Sv/Ev mouse as the control group. **(C)** The path (red) of a representative mouse from each group during the social interaction test. (dotted-circles indicate the interaction zone). **(D)** Distribution of defeat index (DI) scores for defeated and non-defeated 129Sv/Ev mice with each social target. Control/CD1 (n=10), control/129 (n=10), defeat/CD1 (n=20), defeat/129 (n=19). **(E)** Percentage of approaches to the CD-1 that led to avoidance (running away). Control/CD1 (n=10), control/129 (n=10), defeat/CD1 (n=9), defeat/129 (n=10). Data represent mean ± SEM. ****p<0.0001, ***p<0.001, **p<0.01.

### 3.2 Social avoidance is acquired across days of CSDS

During Pavlovian fear conditioning, conditioned fear is acquired as the number of CS-US pairings increase in number. Here, we tested whether social avoidance is similarly acquired across days of CSDS. We tested social interaction repeatedly in mice before CSDS and after 1, 4, 7, and 10 days of defeat, with a novel CD-1 as the social target in each test. To evaluate the effects of stress on bodyweight, defeated and control mice were weighed daily. We found no effect of CSDS on weight change (day 10 weight - baseline weight; t_(13)_ = 0.18, p = 0.86) (Fig. 2B). A two-way repeated measures ANOVA on time with the social target across days revealed no significant stress x day interaction (F_(4,120)_ = 1.194, p=0.32), but a significant effect of day (F_(3.26, 97.85)_ = 7.93, p<0.0001) and stress (F_(1,30)_ = 6.62, p<0.05). Planned group comparisons on each test day revealed that defeated mice interacted with the CD-1 significantly less than the control group after 10 days of CSDS only (p<0.001) (Fig. 2C). These results indicate that 7 days of defeat is not sufficient to induce this behavior and that social avoidance is gradually acquired across days of defeat. To account for the possibility that daily stress exposure in the absence of learning about the CD-1 leads to the emergence of social avoidance, we repeated the experiment in a new cohort of mice using chronic restraint stress. We found that chronic restraint stress induced weight loss (t_(18)_ = 3.57, p<0.01) (Fig. 2D), consistent with previous reports[17, 18]. However, the two-way repeated measures ANOVA on time with the social target revealed no significant group differences (restraint stress x day interaction: F_(4,72)_ = 0.99, p = 0.42; day: restraint stress: F_(1,18)_ = 0.66, p = 0.43), indicating that chronic stress alone is not sufficient to induce social avoidance (Fig. 2E).

**Figure 2.**
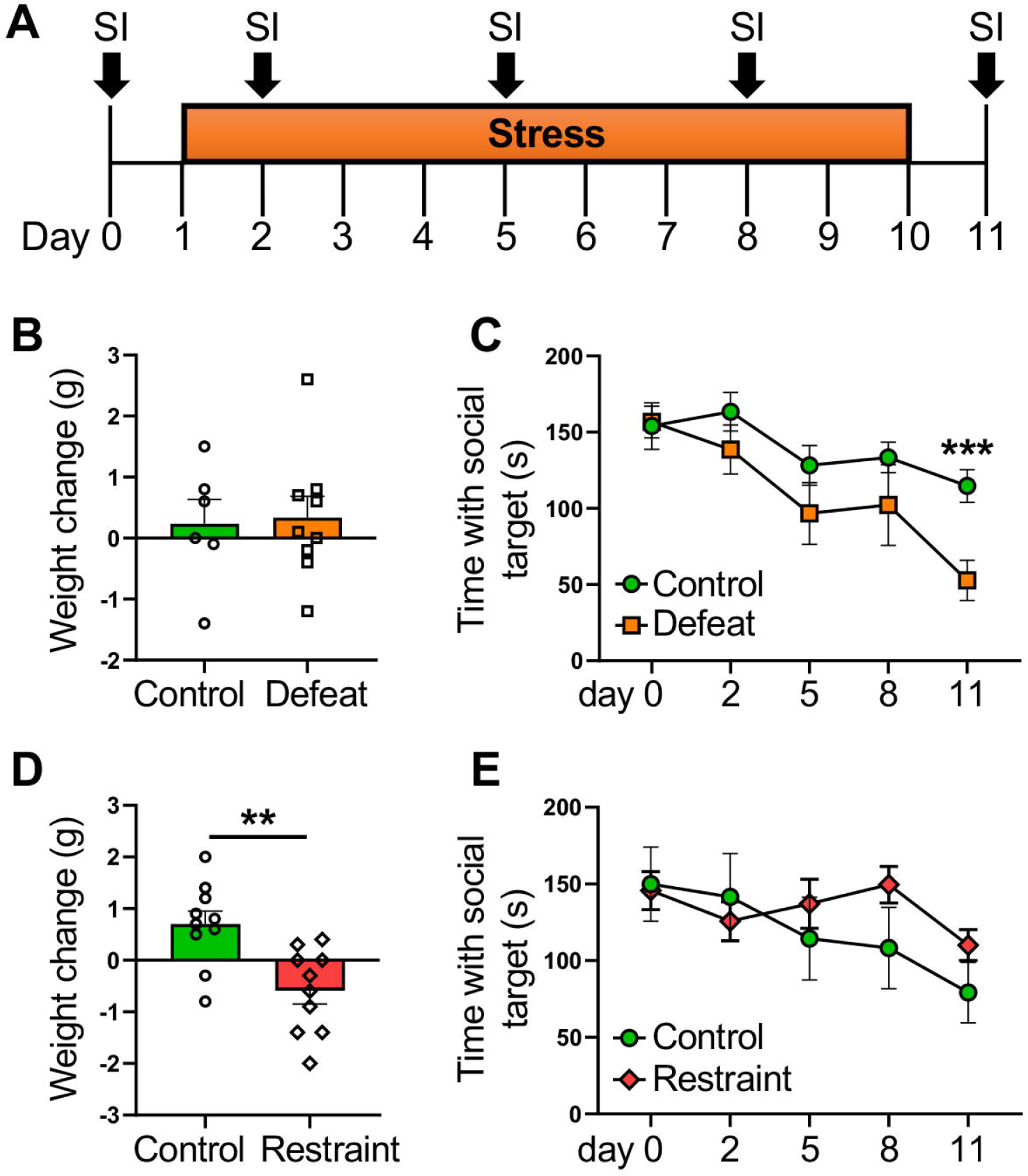
Acquisition of social avoidance is specific to CSDS. **(A)** Schematic of behavioral procedures. Social interaction was tested repeatedly in the same mice with a novel CD-1 as the social target. **(B)** Average change in bodyweight after 10 days of CSDS. Control (n=6), defeat (n=9). **(C)** Time spent with a novel CD-1 across days of defeat (n=16/group). **(D)** Average change in bodyweight after 10 days of restraint stress (1hr/day). **(E)** Time spent with a novel CD-1across days of restraint stress. Unlike CSDS, restraint stress did not lead to the avoidance of a CD-1 (n= 10/group). Data represent mean ± SEM. ***p<0.001 vs. defeated group on day 11, **p< 0.01.

### 3.3 Defeated mice learn to associate white fur with threat

The goal of our next experiment was to identify features of the CD-1 that defeated mice learned to associate with threat during CSDS. We tested whether they learn to fear the white coat of the CD-1, the CD-1 scent, and/or the large body size of the CD-1. Mice were defeated for 10 days and then placed in the social interaction test on the following 2 days, during which one enclosure contained a Swiss Webster mouse, which is visually indistinguishable from the CD-1, or the bedding from a CD-1. The other enclosure remained empty. Then mice were tested with a novel CD-1 or a novel 129Sv/Ev mouse as the social target on the subsequent 2 days. Mice were tested with each target once (4 total tests) and the order of testing was counterbalanced. We separately compared time with each enclosure when the CD-1 was the social target versus each of the other targets (Swiss Webster, CD-1 bedding, 129Sv/Ev mouse). A two-way repeated measures ANOVA on time interacting with the Swiss Webster and the CD-1 revealed a significant main effect of enclosure (F_(1,36)_ = 85.89, p<0.0001), but no significant social target x enclosure interaction (F_(1,36)_ = 3.01, p = 0.09) or main effect of social target (F_(1,36)_ = 0.58, p = 0.45), indicating that defeated mice avoided both the CD-1 and the Swiss Webster to a similar extent. This was confirmed by a planned comparison of time with each social target, which was not significantly different (CD-1 vs. Swiss Webster time, Uncorrected Fisher’s LSD, p = 0.26) (Fig. 3A). In contrast, a two-way repeated measures ANOVA on time interacting with the enclosure containing bedding from a CD-1 and the CD-1 mouse revealed a significant target x enclosure interaction (F_(1, 36)_ = 14.07, p<0.001) with post-hoc tests indicating that mice spent significantly more time with CD-1 bedding than the CD-1 mouse (Uncorrected Fisher’s LSD, p<0.001) (Fig. 3B). Similarly, a repeated measures two-way ANOVA on time interacting with a novel 129Sv/Ev mouse and the CD-1 revealed a significant social target x enclosure interaction (F_(1,36)_ = 8.52, p<0.01). Mice spent significantly more time with the 129Sv/Ev mouse than the CD-1 (Uncorrected Fisher’s LSD, p<0.01) (Fig. 3C), replicating our finding that defeated mice avoid a CD-1 and not a conspecific. To test whether defeated mice learned to fear mice that are bigger than them, we defeated another cohort of mice for 10 days and tested the amount of time they spent interacting with a novel 129Sv/Ev mouse that was ∼35% larger in size. The same mice were then tested with a novel CD-1 as the social target. A two-way repeated measures ANOVA on interaction time revealed a significant social target x enclosure interaction (F_(1,38)_ = 27.53, p<0.0001), with mice spending significantly more time with the larger 129 than the CD-1 (Uncorrected Fisher’s LSD, p<0.0001) (Fig. 3D). Collectively, these results indicate that unlike scent or body size, coat color is a salient feature of the CD-1 that defeated mice learn to associate with threat.

**Figure 3.**
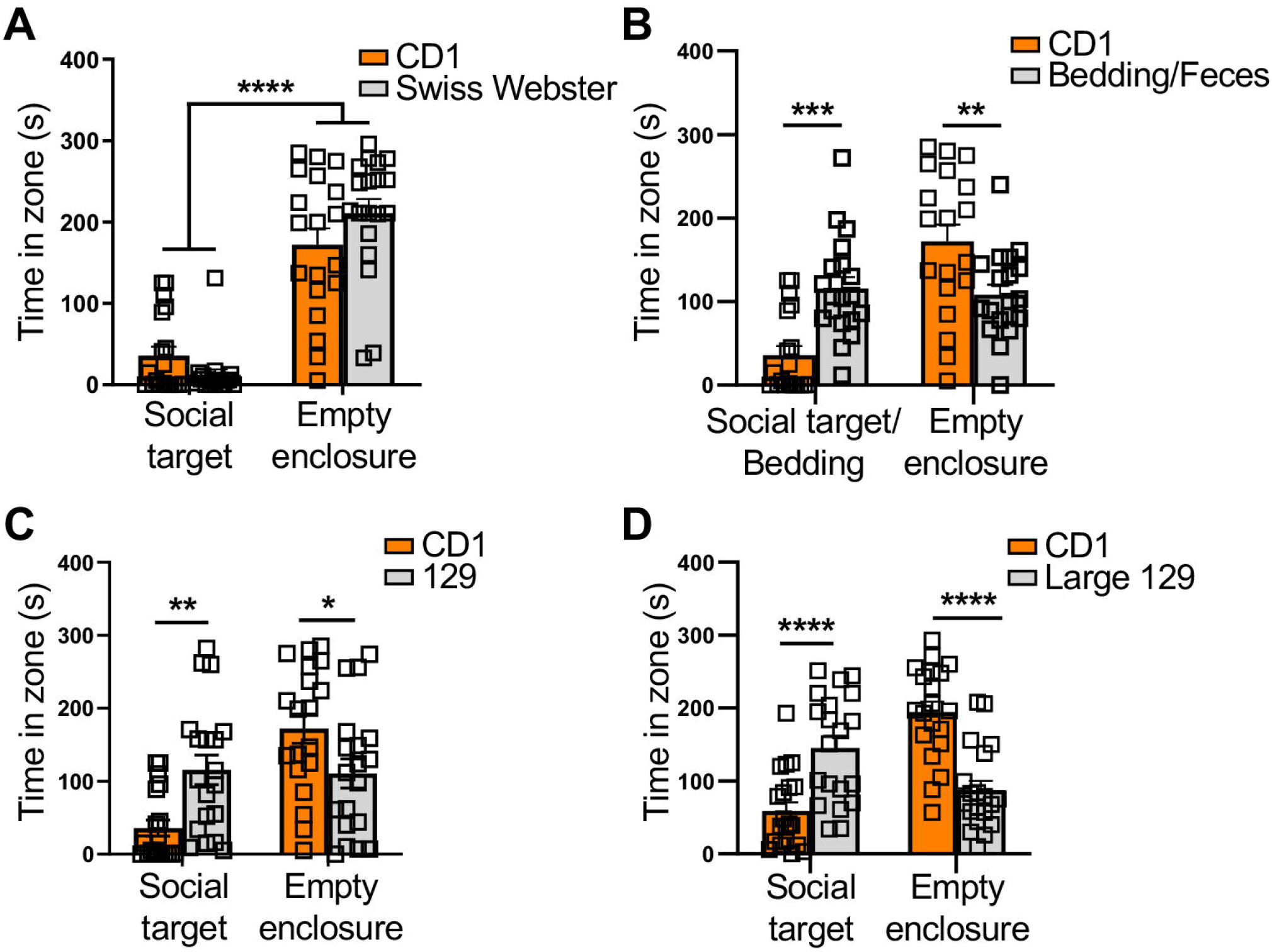
Defeated mice associate white fur with threat. Time defeated mice spent with the empty enclosure and the enclosure containing either a novel CD-1 (n=19), (**A)** a Swiss Webster mouse (n=19), **(B)** bedding containing CD-1 feces (n=19), **(C)** a novel 129Sv/Ev mouse (n=19), or **(D)** a large 129Sv/Ev mouse similar in size to a CD-1 (n=20). Data represent mean ± SEM. *p <0.05, **p < 0.01, ***p < 0.001, ****p < 0.0001.

### 3.4 CSDS-induced social avoidance is subject to extinction

During classical conditioning, extinction occurs when the CS is repeatedly presented in the absence of the US, leading to the cessation of conditioned responding. We hypothesized that if the CD-1 mouse serves as a CS and the attack is a US, then repeated exposure to the CD-1 in the absence of an attack would lead to extinction of social avoidance. We tested this hypothesis by testing defeated mice in the social interaction test 5 times a day for7 consecutive days with either a CD-1 or 129Sv/Ev mouse as the social target (Fig. 4A). There was no physical interaction with either social target during social interaction testing, and defeated mice were therefore never attacked during this time. A two-way repeated measures ANOVA on average time spent interacting with both social targets each day revealed a significant social target x day interaction (F_(6, 102)_ = 2.73, p <0.05), with defeated mice spending significantly less time with the CD-1 than the 129Sv/Ev mouse on the first 4 days of testing only (Day 1, p<0.0001; Day 2, p<0.05; Day 3, p<0.01, Day 4, p<0.05). Post-hoc tests revealed no group differences in interaction time during the last 3 days of testing (Day 5, p = 0.13, Day 6, p = 0.12, Day 7, p = 0.13), indicating that defeated mice gradually stopped avoiding the CD-1 over time (Fig. 4A). This was confirmed with a 2-way ANOVA on time with each social target during the first and last days of testing. There was a significant social target x day interaction (F_(1, 17)_ = 6.80, p< 0.05), with interaction time increasing only when the CD-1 was the social target (Uncorrected Fisher’s LSD, p<0.001) (Fig. 4C). Similarly, a comparison of defeat-index (DI) scores during the first and last days of testing revealed a significant social target x day interaction (F_(1,17)_ = 23.97, p < 0.001), with lower scores on the first than the last day when the CD-1 was the social target (p < 0.0001). In contrast, DI scores remained similarly high when a conspecific was the social target (p = 0.80) (Fig. 4D). These findings indicate that CSDS-induced avoidance of a CD-1 is a learned fear response that can be extinguished.

**Figure 4.**
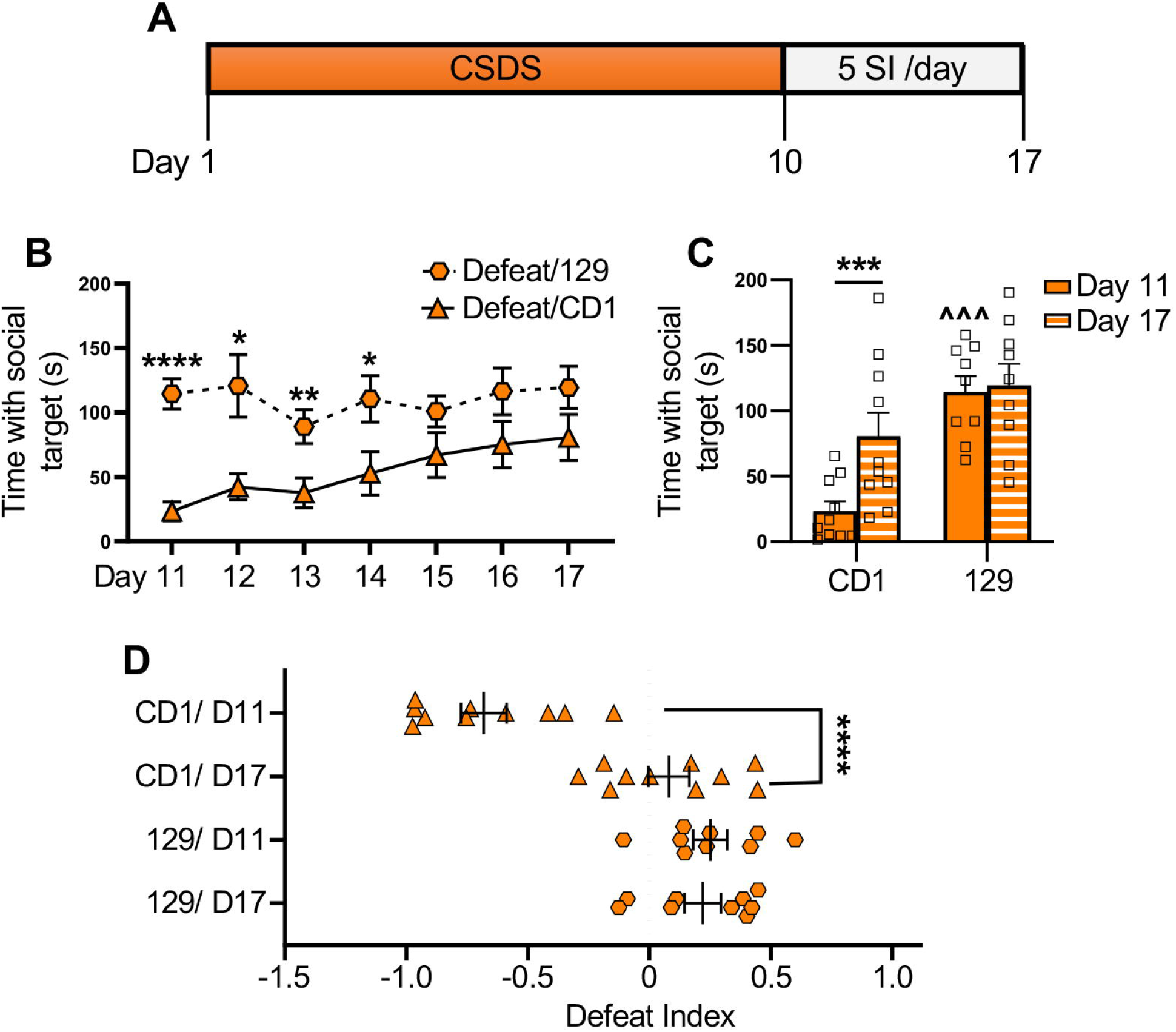
Extinction of social avoidance behavior. **(A)** Schematic of behavioral procedures. Defeated mice were subjected to 5 social interaction tests/day for 7 consecutive days. Mice were either repeatedly exposed to a novel CD-1 mouse or a novel conspecific. **(B)** Average time defeated mice spent with each social target on each day of testing. Data points represent the average of 5 social interaction tests. **(C)** Time spent with each social target on the first and last days of testing. Repeated exposure to the CD-1 in the absence of an attack increased interaction time. **(D)** Distribution of defeat index (DI) scores for defeated mice with each social target on the first (D11) and last days of testing (D17). Data represent mean ± SEM. Defeat/129 (n=9), defeat CD1 (n=10). *p<0.05, **p < 0.01, ***p < 0.001, ****p < 0.0001, ^^^p<0.0001 vs. Day 11/CD-1.

### 3.5 Tone-evoked avoidance behavior after CSDS

If the CD-1 mouse is a conditioned stimulus (CS) and social avoidance is a conditioned response, then pairing the CD-1 with another conditioned stimulus (CS-2) would lead to second-order conditioning, whereby the CS-2 acquires the ability to trigger avoidance behavior. We tested this possibility by pairing a novel CD-1 with a tone (CS-2; 4Khz, 70dB) on each day of CSDS for 10 days and quantifying tone-evoked freezing behavior prior to the appearance of the CD-1. A one-way repeated measures ANOVA on daily freezing revealed a significant effect of day (F_(4.38, 35.07)_ = 5.68. p<0.001), indicating that fear responses to the tone increased as it became associated with the CD-1 (Fig. 5B). Defeated and non-defeated control mice were then placed in the same arena used for the social interaction test, but both enclosures were empty and no social target was present. Instead, interaction with one enclosure triggered the tone (4Khz, 70dB), which played continuously until the mouse left the area (tone-paired enclosure) (Fig. 5A). A two-way ANOVA on time with each enclosure on the day of test revealed a significant main effect of enclosure (F_(1,56)_ = 6.73, p< 0.05) and stress x enclosure interaction (F_(1,56)_ = 10.73, p<0.01). Defeated mice spent less time with the tone-paired enclosure than the other empty enclosure (p<0.001) and less time with the tone-paired enclosure than non-defeated mice (p<0.01) (Fig. 5C), demonstrating avoidance of the enclosure that triggered the tone. These results indicate that a CD-1 mouse is a CS that can be used for second-order conditioning, resulting in fear-induced avoidance behavior in the absence of a social target.

**Figure 5.**
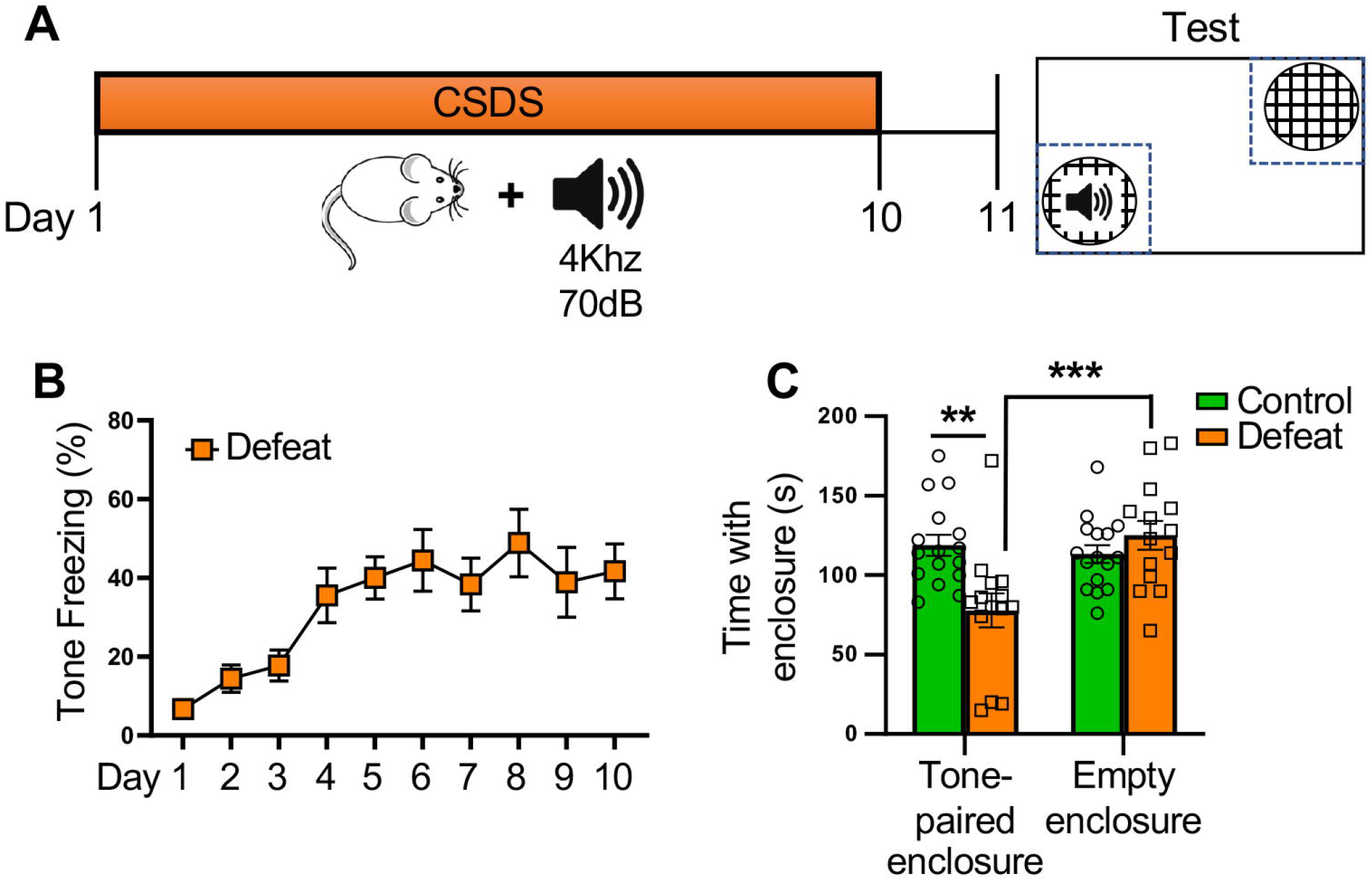
Pairing a CD-1 mouse with a tone leads to second order fear conditioning. **(A)** Schematic of behavioral procedures. On each day of CSDS, defeated mice were allowed to freely explore the home cage of a CD-1 mouse before hearing a tone (30 seconds, 40kHz, 70dB) that was paired with the appearance of the CD-1. On the day of test, mice were exposed to 2 empty enclosures. The tone (40kHz, 70dB) played continuously when mice interacted with one enclosure only (tone-paired enclosure). **(B)** Percentage of time defeated mice spent freezing to the tone prior to the appearance of the CD-1. Mice significantly increased tone-evoked freezing across days of CSDS (n=9). **(C)** Time spent with each enclosure on the day of test. Defeated mice spent significantly less time with the enclosure that triggered the tone (tone-paired) than control mice. Control (n=16), defeat (n=14). Data represent mean ± SEM. **p < 0.01, ***p < 0.001.

### 3.6 Social avoidance of a conspecific correlates with fear generalization

Following fear conditioning, defensive responding not only occurs to stimuli that were explicitly paired with an aversive outcome (CS+), but can generalize to those that were never paired with threat (CS-). Here, we investigated whether this also occurs after CSDS, with fear to the CD-1 (CS+) being generalized to a conspecific, a social target that was never paired with an aversive outcome (CS-). We exposed mice to 10 days of CSDS and tested their social interaction with a CD-1 and a 129Sv/Ev mouse, providing a within-animal measure of avoidance. Then we directly measured fear generalization using a differential fear conditioning paradigm, a commonly used task that tests the ability to discriminate between a tone that was paired with a footshock (CS+) and a different tone that was never paired with a shock (CS-) (Fig. 6A). A two-way repeated measures ANOVA on interaction time revealed a significant main effect of social target (F_(1,38)_ = 4.46, p <0.05) and stress x social target interaction (F_(1,38)_ = 7.08, p<0.05), again confirming that defeated mice avoid a CD-1 and not a conspecific (control/CD-1 vs. defeat/CD-1, p<0.0001; defeat/CD-1 vs. defeat/129, p<0.01) (Fig. 6B). However, interaction time with the 129Sv/Ev mouse was a highly variable response in defeated mice, as indicated by the coefficient of variation, which was significantly higher in defeated than control groups (p<0.001) (Fig. 6C). Our within-animal analysis of DI scores for each social target revealed that 60% of control mice had a stronger preference for the CD-1 than the 129Sv/Ev mouse, which was likely driven by strain novelty (Fig. 6D1). In contrast, 70% of defeated mice had a stronger preference for the conspecific than the CD-1, although the extent of this preference varied (Fig. 6D2). Interestingly, some defeated mice had no preference (20%), a response possibly reflecting generalization of fear to the 129Sv/Ev mouse. Variability in preference for the 129Sv/Ev mouse was also demonstrated by the range of social discrimination scores that we calculated for each defeated mouse (time with 129 – time with CD1) (Fig 6E). We next evaluated avoidance of the conspecific by measuring time with the empty cup when the 129Sv/Ev was the social target and found no difference between groups (t_(38)_ = 0.39, p = 0.70) (Fig. 6F). Interestingly, we did find that 129 avoidance positively correlated with generalization of fear to a non-threatening tone, as measured by freezing to the CS-following differential fear conditioning (r = 0.343, p<0.05) (Fig. 6G). Therefore, the more mice avoided the 129, the more likely they were to be generalizers of fear. Collectively, these results indicate that although CSDS does not decrease average time interacting with a conspecific, there are individual differences in this response that may be attributable to individual differences in generalization of fear to the 129Sv/Ev mouse.

**Figure 6.**
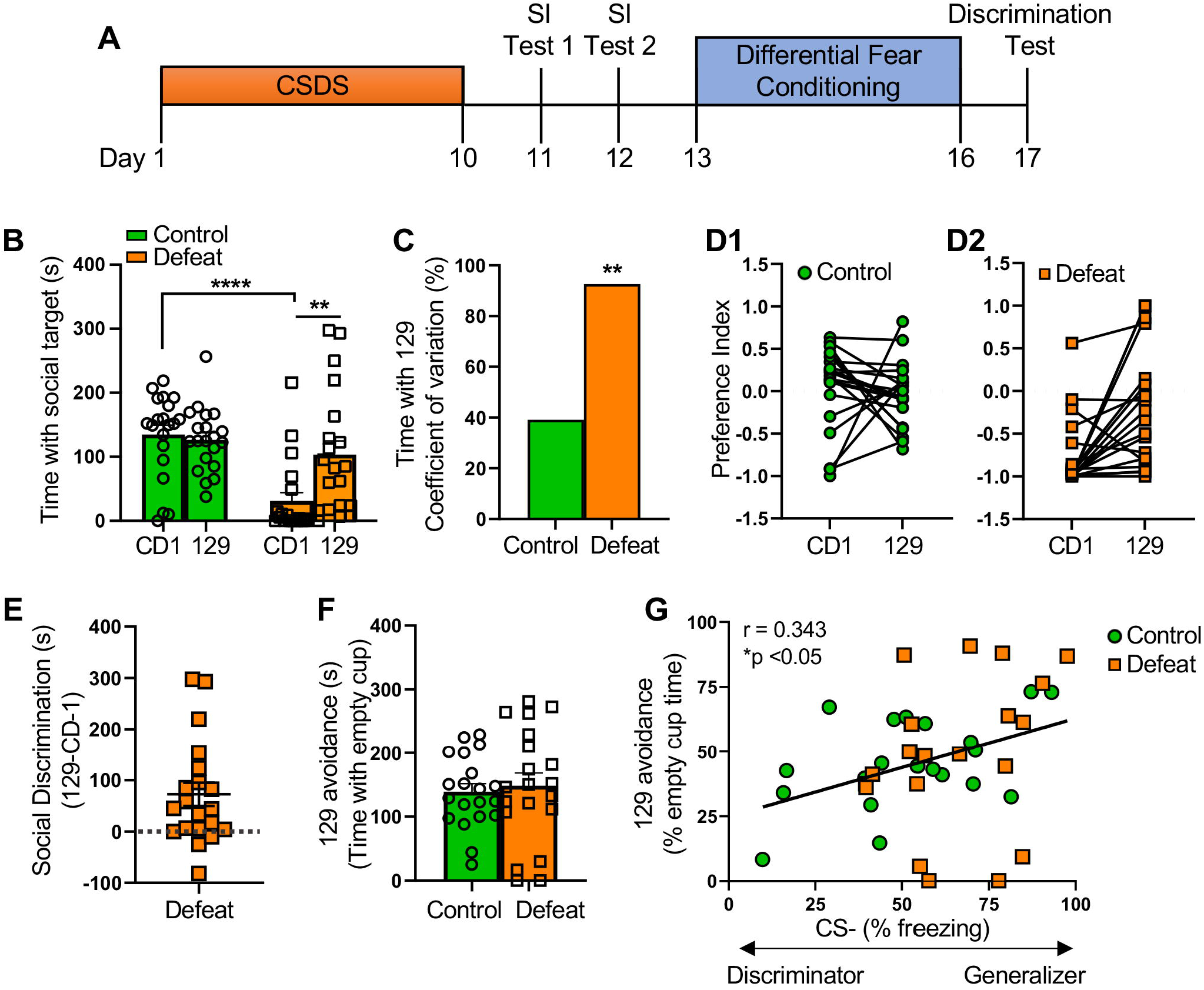
Generalization of fear to a conspecific. **(A)** Schematic of behavioral procedures. After CSDS, mice were tested in 2 social interaction tests with a CD-1 and a conspecific as the social targets. They were then tested in a differential fear conditioning paradigm that evaluated generalization of fear from a CS+ (9kHz tone) to a CS- (2kHz tone). **(B)** Time spent with the enclosure containing each social target. Defeated mice avoided the CD-1 but spent as much time with the 129Sv/Ev mouse as the control group. **(C)** The coefficients of variation of time with the conspecific, which was more variable in the defeated than the control group. **(D)** A preference index is illustrated by graphing the defeat index (DI) with each social target. Lines sloping upward indicate preference for the conspecific. **(E)** Social discrimination in defeated mice calculated as the difference in time spent with each social target. Numbers above zero (dashed line) indicate more time with the 129 than the CD-1, with higher numbers indicating better discrimination between the 2 social targets. **(F)** Time spent with the empty cup when the 129 was the social target. **(G)** Correlation between the percentage of time mice avoid the 129 mouse and percentage of time they freeze to the CS-during the discrimination test. Avoidance of the 129 was associated with generalization of fear to the tone that was never paired with a shock (CS-) (n=20/group). Data represent mean ± SEM. *p<0.05, ** p < 0.01, *** p < 0.001.

## 4. Discussion

CSDS has previously been shown to induce anxiety-like behavior in the elevated plus maze and depressive-like behavior in the sucrose preference test [7]. CSDS-induced avoidance of a CD-1 aggressor has been interpreted as another measure of depressive-like behavior based on the observation that defeated mice also avoid a mouse of the same strain and social avoidance is commonly found in depressed people [6, 8–10]. However, a role for fear learning in CD-1 avoidance is rarely addressed. Here, we show that CSDS leads to avoidance of CD-1, but not a conspecific, indicating that motivation to socialize is intact in the 129Sv/Ev strain. During the social interaction test, defeated mice approach the CD-1 and then run in the opposite direction, activity that is consistent with a threat response. Like associative learning acquired with traditional models of Pavlovian fear conditioning, defeated mice learn to associate the color of the CD-1 coat with threat, leading to gradual acquisition of CD-1 avoidance, a conditioned response that can be extinguished. In addition, pairing a CD-1 with a tone leads to second-order conditioning that results in fear-based avoidance behavior in the absence of a social target, further establishing avoidance behavior as a conditioned response. Finally, social interaction with a conspecific is a highly variable response in defeated mice that may reflect individual differences in generalization of fear to other social targets. Collectively, these findings provide multiple lines of evidence supporting a critical role of fear learning in CSDS-induced avoidance behavior.

We are aware of one other CSDS study conducted in 129Sv/Ev mice that also reported no effect of defeat on avoidance of a conspecific [14]. In contrast, studies conducted in C57BL/6 mice, the most commonly used strain for studying the effects of CSDS, report the opposite[6, 8–10], indicating that CSDS may affect these strains differently. Indeed, we have previously shown that 129Sv/Ev mice are significantly more vulnerable to CSDS than C57BL/6 mice, with only ∼8% of 129s exhibiting resilience compared to ∼67% of C57s [12]. Consequently, it is possible that CSDS has a stronger impact on circuits mediating fear learning in the 129Sv/Ev strain. However, not all CSDS studies with C57BL/6 mice demonstrate avoidance of a conspecific [19, 20]. In addition, some studies show that defeated C57BL/6 mice fail to avoid mice of another strain that either physically resemble them (i.e. Black Swiss mice) [19] or are physically distinct (i.e., 129/Sv) [20, 21], supporting the view that they specifically learned to fear mice that look like the aggressor. The finding that C57BL/6 mice avoid a conspecific after being defeated by other C57BL/6 mice is in line with this interpretation [22]. We demonstrate in our study that interaction with a conspecific is a highly variable response and attribute discrepancies in the literature to this variability. Given that C57BL/6 mice are more stress resilient than 129Sv/Ev mice, and by definition show a wider range of responses to the CD-1, variability in response to a conspecific may also be more pronounced in the C57 strain.

Extinction of social avoidance has been considered previously in studies that re-exposed defeated C57BL/6J mice to the CD-1 by placing them in a familiar aggressor’s home cage (15 minutes/day) for 16 days [20, 21]. During this time, animals were separated by a mesh wall, which is the same way they were housed after being attacked on each day of CSDS, as it provides psychological stress caused by continuous exposure to the CD-1. While this extinction procedure exposes mice to the CD-1 and contextual cues associated with threat (i.e., aggressor’s home cage) in the absence of an attack, it also re-exposes them to psychological stress, potentially minimizing the acquisition of extinction. This may be why only a subset of susceptible mice exhibited extinction, as measured by increases in the social interaction index [21]. In our study, we extinguished animals by repeatedly placing them in an arena that was never associated with threat and measuring daily changes in interaction with the CD-1, allowing us to visualize extinction learning over the course of 7 days of testing. Using this procedure, we found that every defeated animal increased their interaction with the CD-1.

Studies investigating the neural circuits mediating CSDS-induced social avoidance primarily focus on the nucleus accumbens (NAc) and the ventral tegmental area (VTA), areas implicated in depression [7, 8, 23–25]. For example, CSDS increases spine density in both regions [23, 24] and stimulating or inhibiting projections from the VTA to the NAc leads to increased or decreased social interaction in defeated animals [25]. Our study establishing a role for fear learning in social avoidance implicates the involvement of the basolateral nucleus of the amygdala (BLA), a stress sensitive region [26–28] known to be essential for fear learning [29] and learned avoidance behavior [30–32]. Glutamatergic projections from the BLA to the NAc synapse on the same medium spiny neurons that receive input from the mesolimbic dopamine system [33], indicating that the BLA is anatomically positioned to modulate these circuits that are already known to mediate social avoidance. The involvement of the BLA is further implicated in a recent CSDS study demonstrating that there are differentially expressed genes in the BLA of resilient and a subgroup of susceptible mice [21]. Understanding how amygdala-based fear circuits might interact with reward circuits to promote CSDS-induced changes in social interaction would be important in future studies.

## 5. Conclusions

CSDS-induced avoidance of a CD-1 aggressor is often interpreted as depressive-like behavior. Using 129Sv/Ev mice, a strain that is particularly vulnerable to stress, we show that associative learning occurs during each defeat session and that avoidance behavior is a learned response to threat. Our findings indicate that CSDS is a paradigm that can be used for studying fear learning involving a social cue, a process that occurs in humans experiencing bullying. Future CSDS studies identifying how fear circuits and reward circuits interact to promote social avoidance may provide insight into novel treatment options for individuals affected by this type of social stress.

## Acknowledgments

We thank members of the animal care staff at Hunter College, particularly Barbara Wolin and Sonia Acevedo. This project was supported by NIMH R21MH114182 (N.S.B. & E.L.), NIMH R21MH135430 (N.S.B. & E.L.), NIMH R01MH118441 (E.L.), & PSC-CUNY Awards (N.S.B).

## Conflict of Interest Statement

The authors declare that they have no conflict of interest.

## Notes

### Competing Interest Statement

The authors have declared no competing interest.

